# SMAD4 promotes formation of terminally differentiated CTLs that localize in the microvasculature of the lungs but are excluded from the lumen of the airways

**DOI:** 10.1101/2023.11.16.567437

**Authors:** Jenny Suarez-Ramirez, Karthik Chandiran, Linda S Cauley

**Affiliations:** CaroGen Corporation Farmington, CT. USA; School of Biology, Indian Institute of Science Education and Research, Thiruvananthapuram, Kerala, India; Department of Immunology, UCONN Health, Farmington, CT. USA

**Author notes:** **CORRESPONDENCE:** Dr. Jenny Suarez-Ramirez.

## Abstract

Cytotoxic T lymphocytes (CTLs) circulate around the body searching for infected and transformed cells, that undergo apoptosis when lytic granules are delivered into the cytoplasm. To find pathogens that propagate in different tissues, naïve CD8 T cells differentiate into heterogeneous populations of effector (T_EFF_) and memory CD8 T cells with different migratory properties. Several subsets can be identified using antibodies that recognize surface receptors that are expressed at specific stages during CD8 T cell differentiation. Although flow cytometry is a powerful method for tracking antigen specific CTLs during a dynamic immune response, the data provide little information about the distribution of cells in specific anatomical compartments. In this study, confocal imaging was used to explore how signaling via SMAD4 influenced the tissue-tropism of antigen specific CTLs during respiratory infection. During microbial infection, wildtype CTLs gave rise to terminally differentiated T_EFF_ cells that expressed KLRG1 and CX_3_CR1 at high levels and localized in the microvasculature of the lungs. However, both markers were expressed at reduced levels on SMAD4-deficient CTLs, which preferentially entered the lumen of the airways. These disparate homing properties emphasize the important contributions of SMAD signaling pathways to cell-mediated immunity.

## Introduction

During respiratory infection, epithelial cells are important sites of viral replication. Infected cells are eliminated by CD8 T cells that undergo clonal expansion in secondary lymphoid organs (SLO) and disperse to the site of infection. Migrating CTLs leave the circulation by crossing the endothelial barrier at exit points in venules. Most T_EFF_ cells disappear as the infection is cleared, while specialized memory CD8 T cells provide sustained immune surveillance against reinfection. Three subsets of memory CD8 T cells can be distinguished based on their migratory properties. Central memory CD8 T cells (T_CM_) use blood and lymph to circulate through SLO, while effector memory CD8 T cells (T_EM_) lack lymphoid homing receptors and survey peripheral tissues(1). Members of the third subset are non-circulating tissue resident memory CD8 T cells (T_RM_) that leave the circulation and become lodged in the local tissues(2, 3). T_RM_ cells are poised to augment immunity by intercepting new pathogens at the point of entry into the body(4). The ratios of T_EFF_ and long-lived memory CD8 T cells vary during infections with different types of pathogens, emphasizing the influence of inflammatory mediators during T cell differentiation (5).

The airways are lined with pseudostratified epithelial cells, that are connected by protein structures known as tight junctions (TJs) and adherens junctions (AJs)(6). At AJs cellular interactions are mediated by calcium-dependent adhesion molecules (Cadherins) that help maintain tissue integrity (6, 7). Transforming growth factor β (TGFβ) is a pleiotropic cytokine that plays a key role wound healing and tissue remodeling by inducing epithelial to mesenchymal transition (EMT) and angiogenesis (8, 9). During EMT, stromal cells acquire migratory properties as epithelial (E)-Cadherin is down regulated by TGFβ, while neural (N)-cadherin is induced(10, 11). Both cadherins provide binding sites for adhesion molecules that are expressed on different subsets of antigen experienced CTLs (12, 13). Killer cell lectin-like receptor G1 (KLRG1) is an inhibitory receptor with a tyrosine-based inhibitory motif in the cytoplasmic domain and is expressed on terminally differentiated T_EFF_ cells(14, 15), whereas CD103 (αEβ7 integrin) is expressed on naïve CD8 T cells and large numbers of noncirculating T_RM_ cells (T_RM_)(3, 16). Regulation by TGFβ prevents dual KLRG1 and CD103 expression on the same CTLs(17). CD103 is induced by constant stimulation with TGFβ(18), whereas KLRG1 is down-regulated(19). Cytokines that induce KLRG1 expression on CTLs *in vitro* have not been identified (20).

Canonical signals for members of the transforming growth factor family are mediated a network of structurally related molecules known as SMAD proteins(21). The SMAD signaling cascade includes an adaptor protein (SMAD4) that shuttles complexes of phosphorylated R-SMADs into the nucleus for gene regulation. We previously used Cre-lox recombination to prevent SMAD4 expression in CD8 T cells and analyzed homing receptor expression during microbial infection. After SMAD4 was ablated abnormally large numbers of CTLs expressed CD103 in the spleen and lungs, while very few CTLs expressed KLRG1 or CD62L(22). These altered phenotypes revealed defects during formation of terminally differentiated T_EFF_ cells and T_CM_ cells(22, 23). We also used transcriptome analysis to monitor changes in gene expression after TCR stimulation. Our data showed that TGFβ and SMAD4 are essential components of two distinct signaling pathways that cooperatively regulate homing receptor expression by altering the expression levels of the same genes in opposite directions(17). Importantly, signaling via SMAD4 altered the migratory properties of newly activated CTLs via a mechanism that did not involve TGFβ(17, 24).

Flow cytometry is often used track CTL responses during microbial infection but cannot provide precise information about the distribution of cells within individual tissues. To further examine the role of the SMAD signaling cascade during T cell migration, we used confocal microscopy to visualize pathogen specific CTLs in the lungs during respiratory infection. After transfer, wildtype donor cells gave rise to KLRG1^+^ T_EFF_ cells that localized in blood vessels around the alveoli, whereas SMAD4-deficient CTLs mostly lacked KLRG1 expression and preferentially entered the lumen of the airways. Further work may reveal how the adhesive properties of cadherin-binding proteins influence tissue localization.

## Results

To track antigen specific CTLs during infection, mice were infected with recombinant pathogens that express the SIINFEKL peptide derived from chicken ovalbumin(25). Two pathogens were used for this study: a recombinant strain of influenza A virus (X31-OVA) that encodes SIINFEKL in the neuraminidase stalk (26) and *Listeria monocytogenes* which encodes a truncated form of ovalbumin (LM-OVA)(27). To obtain a supply of donor cells for transfer studies, we used OTI mice with CD8 T cells that express a transgenic T cell receptor (TcR) that is specific for the SIINFEKL peptide(25). To study SMAD dependent signaling pathways in pathogen-specific CTLs, we used distal Lck-Cre for targeted gene ablation (28). Since DNA-rearrangement occurs after positive selection of thymocytes, loss of signaling via TGF-β receptor II or SMAD4 has minimal impact on the functional properties of naïve CD8 T cells(17, 22). The OTI mice were bred with mice that lack TGFβ receptor II (OTI-TR2KO) or SMAD4 (OTI-S4KO).

Terminally differentiated T_EFF_ cells express the fractalkine receptor (CX_3_CR1) at KLRG1 high levels(5, 17). To visualize these CTLs by confocal microscopy, we bred OTI-Ctrl and OTI-S4KO mice with reporter mice that express green fluorescent protein (GFP) under the control of the CX_3_CR1 promoter(29). CD45.1^+^ donor cells were transferred to C57/BL6 mice before infection with X31-OVA. On different days post infection (dpi), fragments of fixed lung tissue were stained with antibodies to visualize the donor cells (CD45.1), airways (EpCAM) and blood vessels (CD31) **(Fig 1**). Images were recorded at 20X normal magnification.

**Figure 1.**
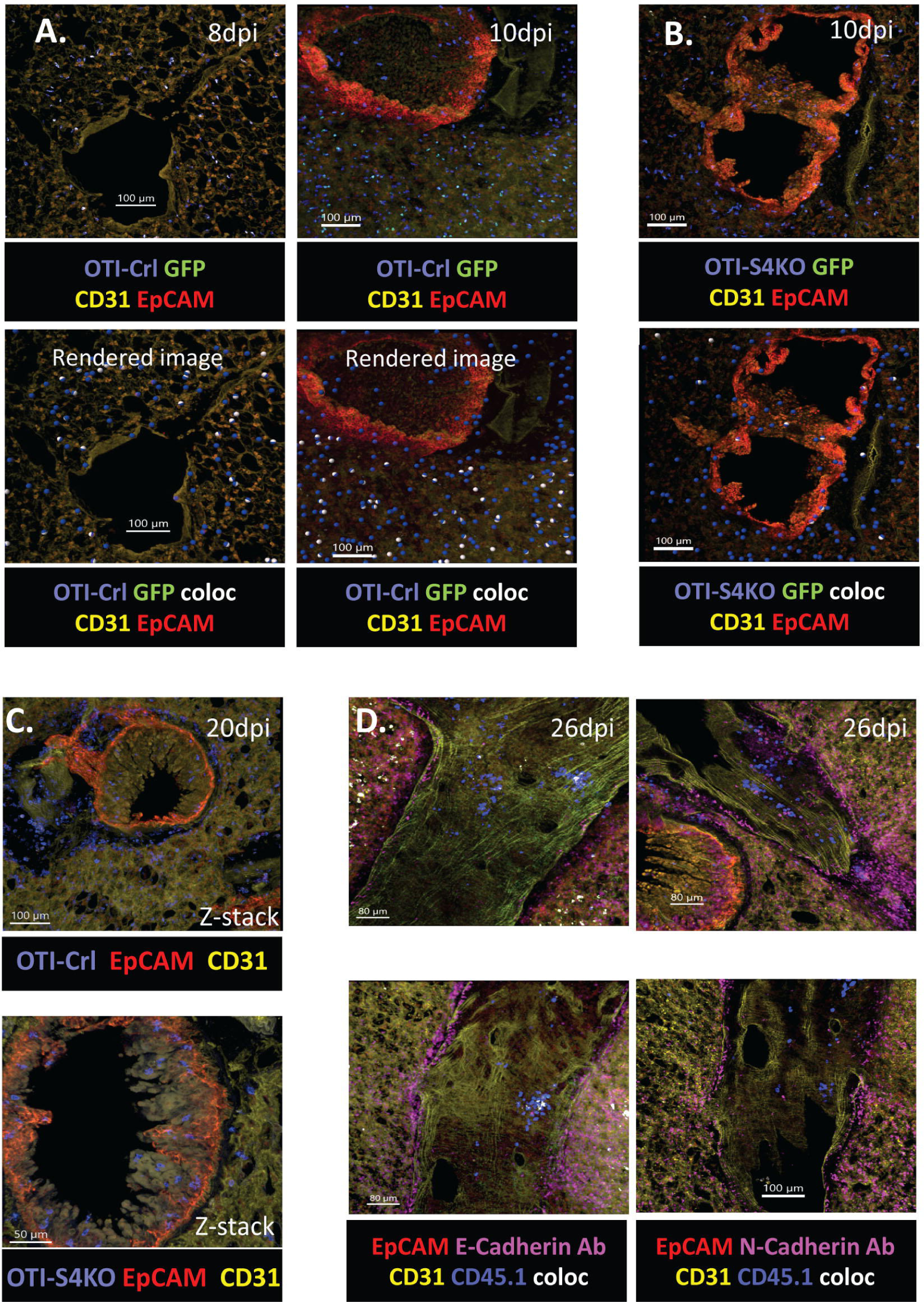
Wildtype CTLs express CX3CR1 at high levels in the microvasculature of the lungs. GFP reporter mice were used to analyze CX3CR1 expression on donor cells in the lungs. OTI-Ctrl and OTI-S4KO cells were transferred to B6 mice before infection with X31-OVA. Tissues were analyzed at 8-10dpi **A)** lmaris colocalization function was used to identify CD45.1+ cells that expressed GFP. OTI-Ctrl cells expressed GFP at high levels in the microvasculature of the lungs. **B)** Most OTI-S4KO cells expressed GFP at low levels. **C)** OTI-S4KO cells entered the the airways 20dpi. **D)** Clusters of OTI-Ctrl cells were attached to the walls of large blood vessels at 26dpi.

At 10dpi, the microvasculature of the lungs was populated with OTI-Ctrl cells that expressed GFP at high levels **(Fig. 1A)**. We used the Imaris® colocalization function to highlight CD45.1^+^ cells that expressed GFP (Rendered images). The microvasculature was also populated with some OTI-S4KO cells that expressed GFP at low levels **(Fig. 1B)**. At 20dpi, OTI-Ctrl cells were present in the peribronchial space **(Fig. 1C -top)** and OTI-S4KO cells embedded in the walls of a large airway **(Fig. 1C -bottom.)**

To examine the distribution of T_EFF_ cells in more detail, we used LM-OVA to generate large numbers of CTLs that expressed KLRG1(30). Mixed OTI-Ctrl and OTI-S4KO cells (1:1 ratio) were transferred to C57BL/6 mice 48hrs before infection. On the days shown, flow cytometry was used to measure the ratios of donor cells in BAL fluid (B) Lungs (L), mediastinal (M) cervical (C) inguinal (I) lymph nodes and spleens (S) **(Fig. 2A).** The gray bars indicate numbers of donor cells that expressed KLRG1 and/or CD103 **(Fig. 2B).** At 8dpi the lungs contained substantial numbers of OTI-Ctrl cells that expressed KLRG1 at highly levels. However KLRG1 was not expressed on OTI-Ctrl cells that entered the airways (BAL fluid). Conversely, OTI-S4KO cells preferentially entered the lumen of the airways (blue bars) and mostly lacked KLRG1 expression. We previously used intravascular (i.v.) staining to analyze anti-viral CTLs at 40dpi with IAV and found that virtually all KLRG1^+^ CTLs were located inside the blood vessels(17, 22). Taken together, these studies suggest that signaling via SMAD4 impedes migration of terminally differentiated CTLs from the vasculature into the airways.

**Figure 2.**
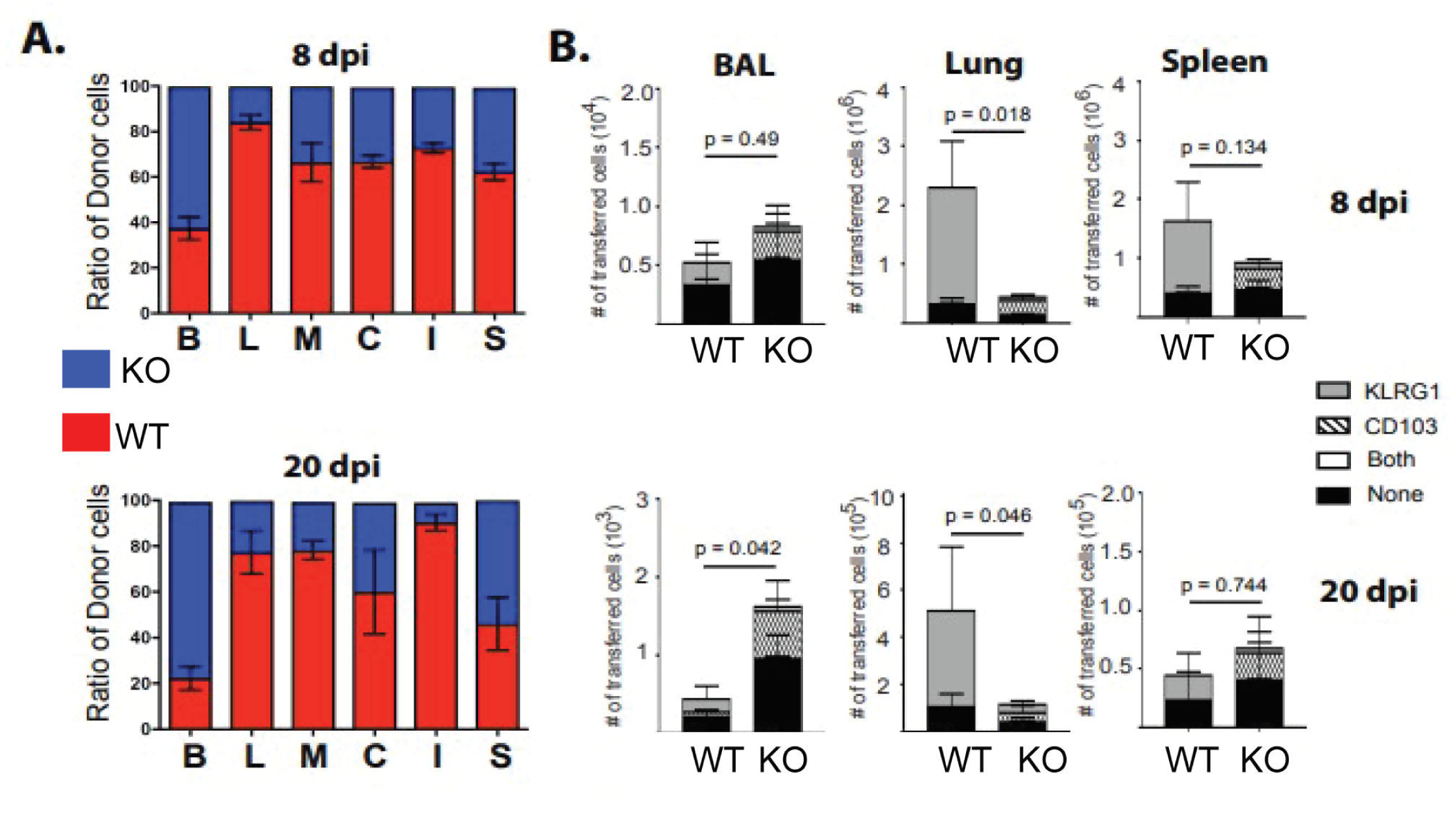
OTI-S4KO cells preferentially enter the lumen of the airways. Mixed donor cells were transferred to C57BL/6 mice before infection with LM-OVA. Tissues were harvested on the days shown and donor cells were analyzed by flow cytometry. Data are mean+ SD n = 5/group. **A)** Colored bars show ratios of OTI-Ctrl (red) and OTI-S4KO (blue) in BAL fluid (B), lung parenchyma (L), MLN (M), CLN (C), ILN (I) and spleen (S). **B)** Numbers of OTI-Ctrl (WT) and OTI-S4KO (KO) cells that expressed KLRG1 and/or CD103 in the lungs and spleen.

The vessels that carry blood and lymph around the body are lined with a continuous layer of endothelial cells, that create a barrier between the circulation and peripheral tissues. When leukocytes interact the endothelial cells, tyrosine phosphorylation of VE-cadherin destabilizes of the endothelial junction to permit diapedesis(31). To visualize OTI-Ctrl and OTI-S4KO cells in the lungs, naïve donor cells were transferred to C57BL/6 mice before infection with LM-OVA. Fixed lung tissue was imaged at 10dpi **(Fig. 3)**. The images show OTI-Ctrl cells interacting with the basement membrane around a large airway **(Fig. 3A)**, whereas OTI-S4KO transversed the epithelial layer **(Fig. 3B)**. Some OTI-Ctrl cells were attached to walls of large blood vessels **(Fig. 3C),** whereas similar vessels contained very few OTI-S4KO cells **(Fig. 3D)**. Other images that were taken after H&E staining also show lymphocytes attached to the walls of some blood vessels **(Fig. 3E)**.

**Figure 3.**
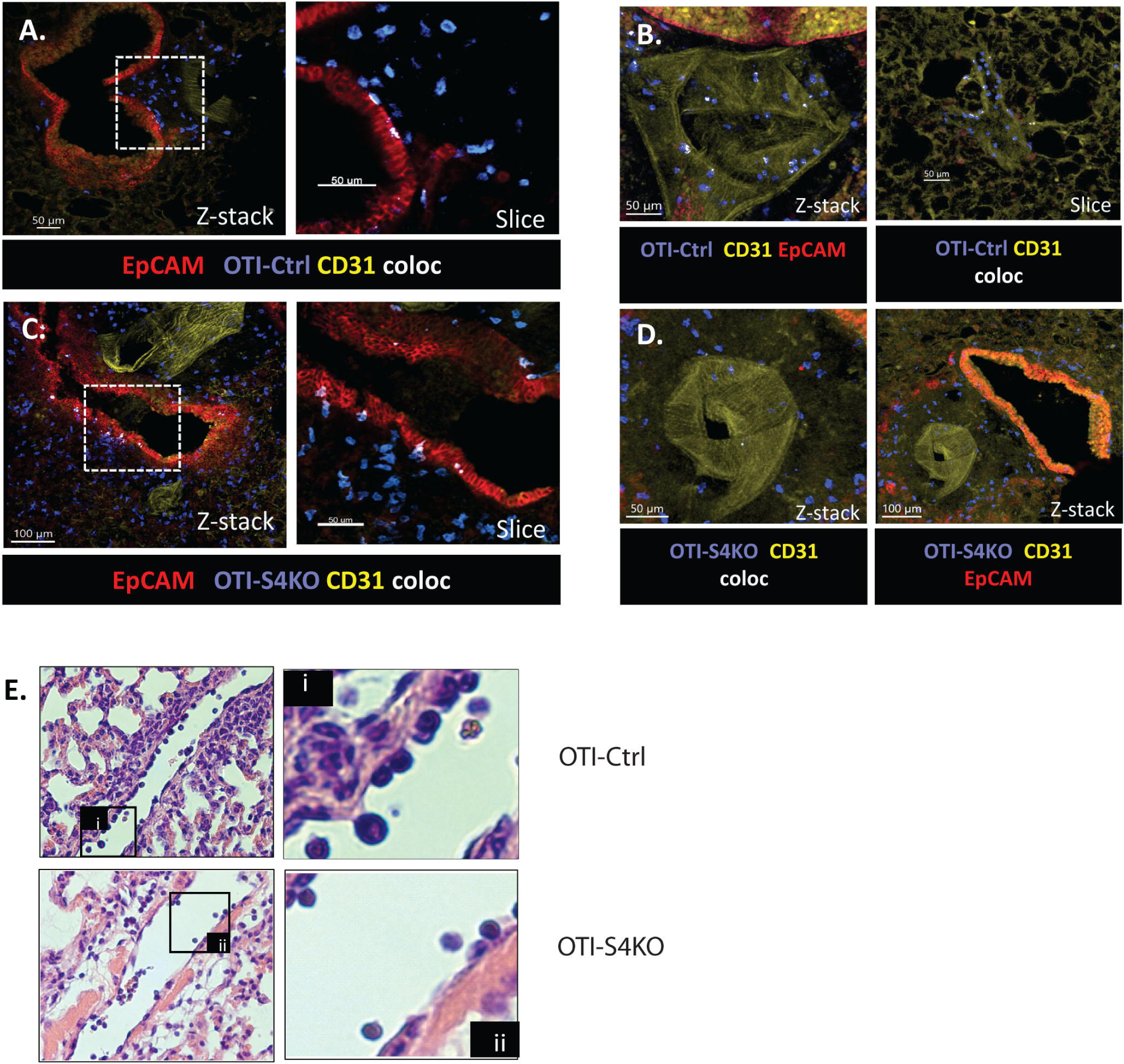
OTI-S4KO cells enter the airways after infection with LM-OVA. CD45.1+ donor cells were transferred to C57BL/6 mice before infection with LM-OVA. At 10dpi, fixed lung tissue was stained with antibodies that are specific for CD45.1 to detect donor cells (blue), EpCAM (red) to detect large airways and CD31 for blood vessels (yellow). **A)** OTI-Ctrl cells interact with the basement membrane around an airway. The boxed regions indicate the positions of enlarged images. **B)** OTI-Ctrl cells were attached to the walls of a large blood vessel. **C)** OTI-S4KO cells traverse the epithelial barrier during transit into the airway. **D)** Very few OTI-S4KO cells were attached to the blood vessel wall. **E)** H&E staining shows lymphocytes attached to the walls of blood vessels in the lungs. The boxed regions indicate

KLRG1 and CD103 are cadherin-binding proteins (32, 33). The extracellular domain of KLRG1 binds to three members of the cadherin family, including E-cadherin which expressed on stromal cells around the airways (32). We used confocal microscopy to explore whether antigen specific CTLs interact with cadherin^+^ cells in the lungs. To generate KLRG1^+^ T_EFF_ cells, mixed OTI-Ctrl and OTI-S4KO cells (1:1 ratio) were transferred to C57BL/6 mice before infection with LM-OVA. At 8 &12dpi, fragments of fixed lung tissue were stained with antibodies specific for CD45.1 (donor cells), CD31 (yellow), EpCAM (red) and E-cadherin (magenta) **(Fig. 4)**. Interactions between CD45.1^+^ cells and E-cadherin were highlighted using the Imaris® colocalization function (rendered images). We found OTI-S4KO cells (blue beads) interacting with E-cadherin in the peribronchial and perivascular space, whereas similar interactions between OTI-Ctrl cells (green beads) and E-cadherin were less obvious.

**Figure 4.**
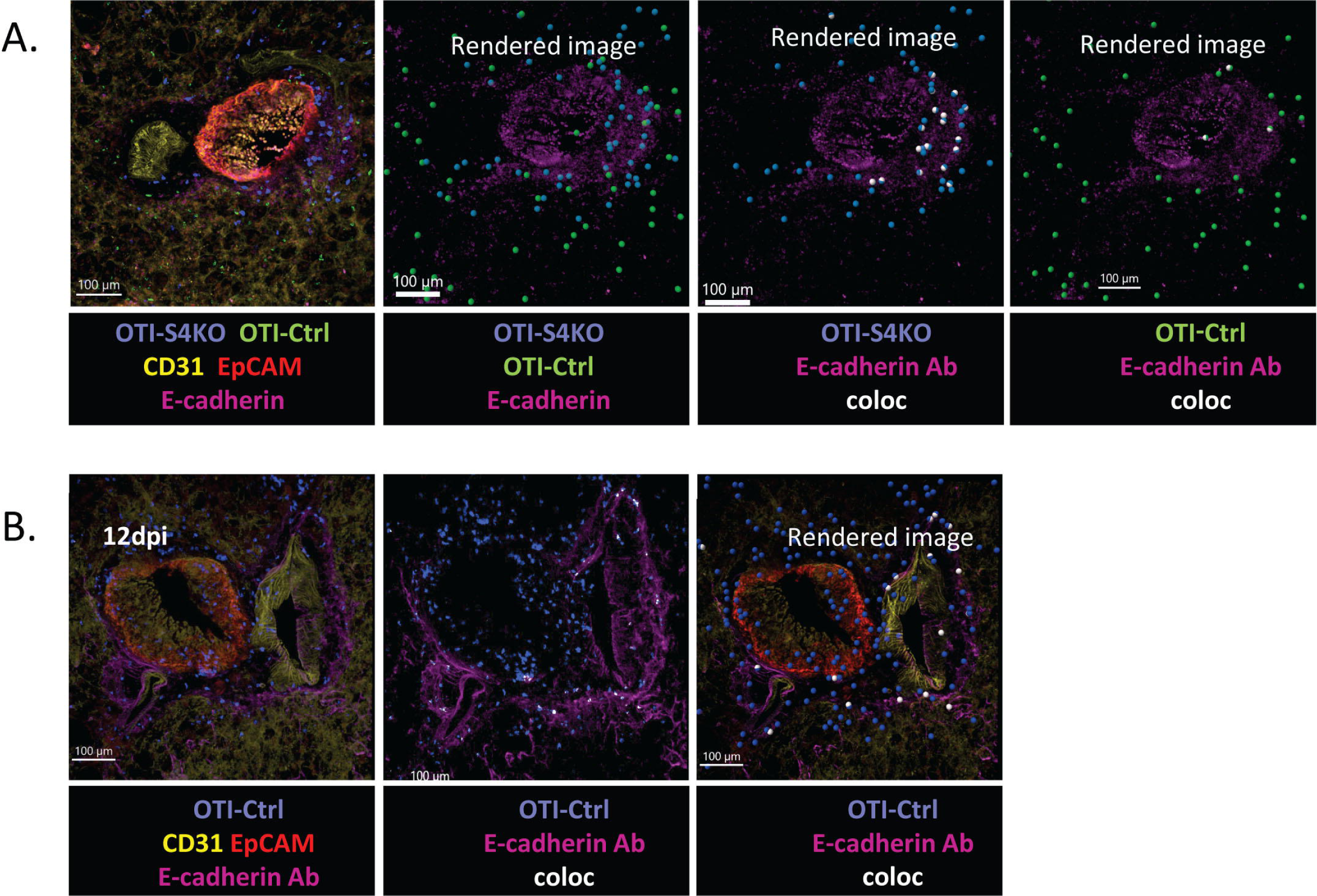
OTI-S4KO cells colocalize with E-cadherin in the peri bronchial space. **A).** Mixed OTI-Ctrl (green) and OTI-S4KO cells (blue) were transferred to C57BL/6 mice before infection with LM-OVA. Fixed lung tissue was stained and imaged 8dpi. Contacts between CD45.1+ donor cells (blue) and E-cadherin (magenta) are shown in white. **B).** Mixed OTI-Ctrl were transferred to B6 mice before infection with X31-OVA. The lungs were imaged 12dpi.

N-cadherin is an alternative ligand for KLRG1 and is expressed on a variety of mural cells and pericytes(10, 34). We used two different strains of reporter mice to visualize N-cadherin^+^ mural cells in the lungs. mTmG dual reporter mice express membrane-bound red fluorescent protein (mRFP) in all cells(35). When Cre recombinase is expressed, mRFP is replaced with membrane-bound green fluorescent protein (mGFP). We crossed mTmG mice with mice that express tamoxifen inducible Cre (ERT-Cre) under the control of the N-cadherin promoter(36). The progeny are called Ncad-mGFP mice. For other experiments, N-cadherin ERT-Cre mice were crossed with mice that express soluble dtomato RFP (37). These mice are called Ncad-dTom.

To visualize interactions between antigen-specific CTLs and N-cadherin^+^ mural cells, we transferred mixed OTI-Ctrl and OTI-TR2KO cells (1:1 ratio) to Ncad-mGFP mice before infection with X31-OVA. The recipient mice were injected twice with tamoxifen to induce Cre expression. Flow cytometry was used to analyze OTI-Ctrl and OTI-TR2KO cells for KLRG1 expression **(Fig 5A)**. The images showed GFP^+^ mural cells surrounding discrete sections of large blood vessel in the lungs **(Fig 5B).** To detect protein expression, we stained lung tissue from Ncad-mGFP mice with N-cadherin antibodies **(Fig 5C).** The antibody staining was not entirely consistent with the GFP reporter, indicating the presence of some soluble N-cadherin attached to extracellular matrix(38). We used the Imaris® colocalization function to detect interactions between GFP^+^ mural cells (green) and CD45.1^+^ donor cells (magenta) **(Fig 5D).** Imaris® was also used to distinguish OTI-Ctrl from OTI-TR2KO cells **(Fig 5E)**. A side by side comparison showed some OTI-TR2KO cells (blue beads) concentrated around a large blood vessel (marked region), whereas OTI-Ctrl cells (yellow beads) were primarily in the microvasculature.

**Figure 5.**
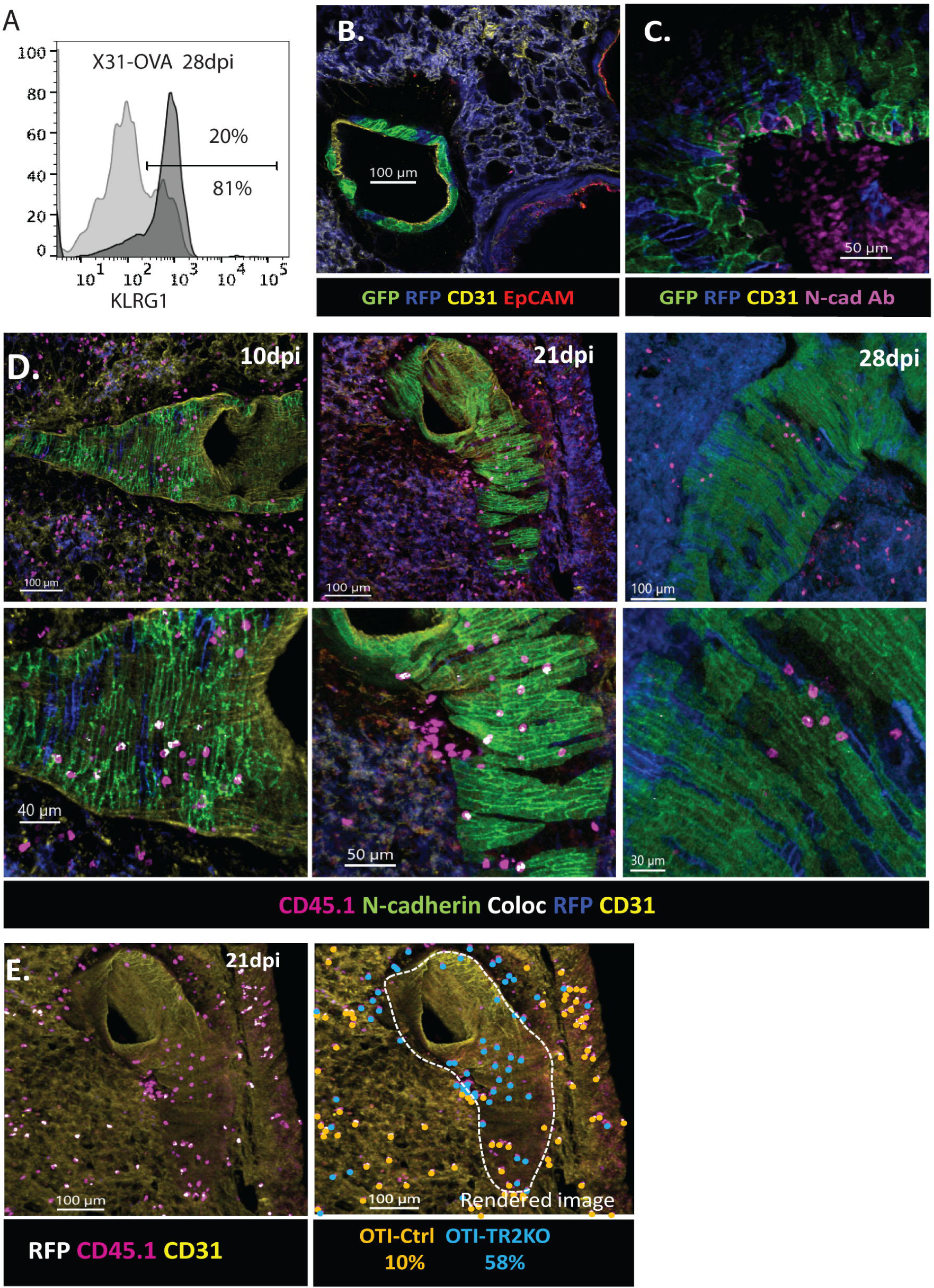
Ncad-mGFP reporter mice were used to visualize N-cadherin+ mural cells (GFP) in the lungs. A) Overlaid histograms show KLRG1 expression on OTI-Ctrl (Light gray) and OTI-TR2KO cells (dark gray). B) Cross section of a blood vessel surrounded by N-cadherin+ mural cells. C) Antibodies were used to stain N-cadherin in the lungs (Magenta). D&E). Mixed OTI-Ctrl & OTI-TR2KO cells were transferred to Ncad-mGFP reporter mice before infection with X31-0VA. Lungs were imaged on different dpi. D) The lmaris colocalization function was used to detect interactions between CD45.1+ donor cells (magenta) and GFP+ mural cells (green). E) Rendered images show the percentages of donor cells that are located outside the marked region (white line).

We were concerned that some GFP^+^ cells may have been rejected from the tissues of Ncad-mGFP mice, so we transferred other OTI-TR2KO cells to Ncad-dTom mice and imaged the lungs at 17dpi with X31-OVA. Consistent with the Ncad-mGFP reporter, discrete sections of some large blood vessels were surrounded with dTom+ mural cells **(Fig 6A)**. Imaris was used to detect interactions between OTI-TR2KO cells (magenta) and RFP+ mural cells (Blue) along the vessel wall. We also imaged MLNs from Ncad-dTom mice at 26dpi and found RFP^+^ cells surrounding the HEVs **(Figs 6C & 6D)**.

**Figure 6.**
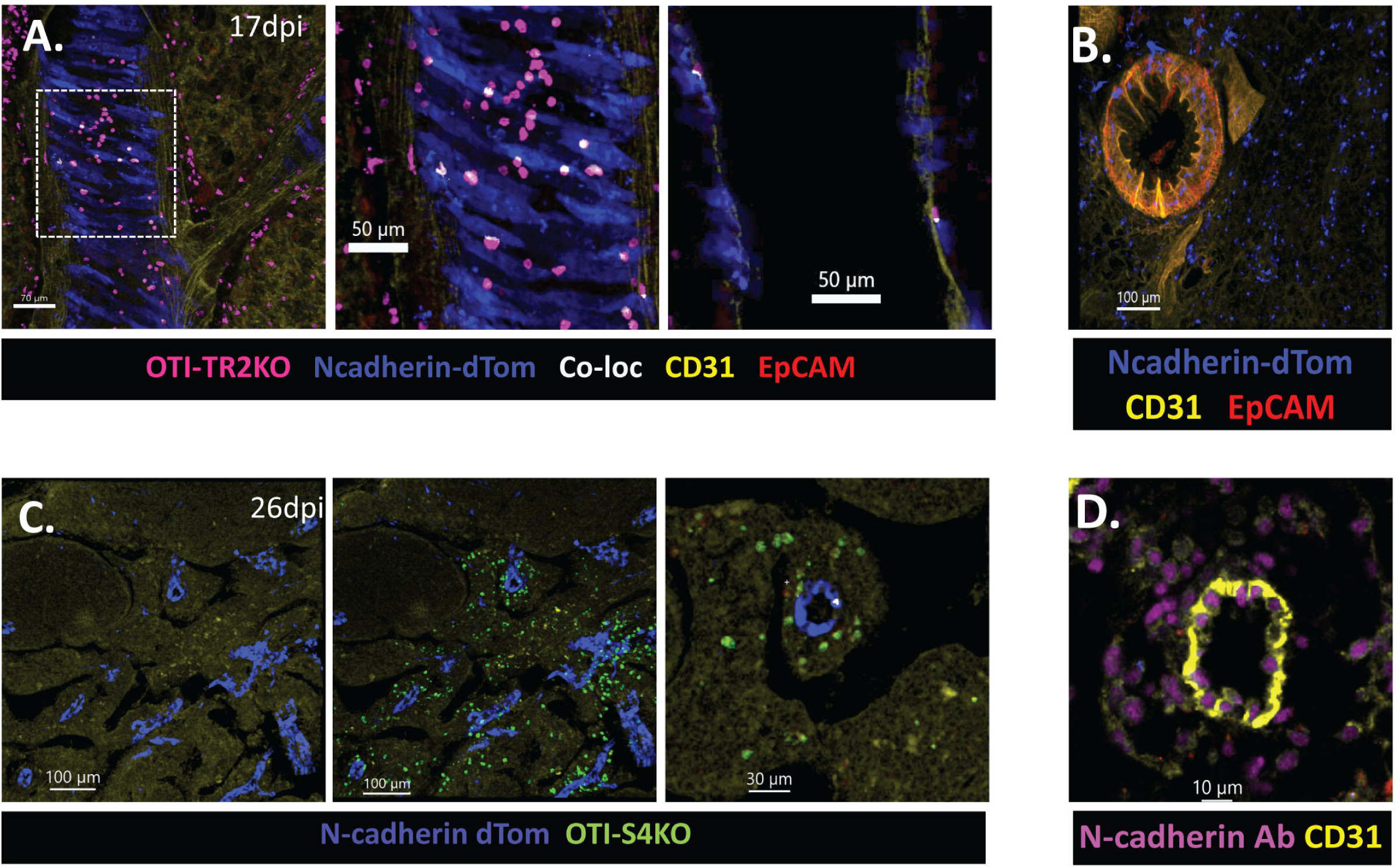
Ncad-dTom reporter mice were used to visualize N-cadherin+ cells in the lungs and MLNs. OTI-TR2KO cells were transferred to Ncad-dTom reporter mice before infection with X31-0VA. The lungs were imaged at 17dpi **A)** N-cadherin+ mural cells (blue) surround a large blood vessel in the lungs. Interactions between OTI-TR2KO cells (magenta) and N-cadherin+ cells are shown in white. The white box indicates the position of the enlarged image. **B)** N-cadherin+ cells in the lung parenchyma. **C)** MLNs from Ncad-dTOM mice were imaged 26dpi. HEVs express N-cadherin (blue). **D)** Crossection of a HEV in the MLN from C57BL/6 mouse. MLN was stained with antibodies specific for CD31 (yellow) and Ncadherin (magenta).

We used mTmG reporter mice to generate OTI-S4KO cells that expressed mGFP for dual transfer experiments. Mixed OTI-TR2KO and OTI-S4KO cells (1:1 ratio) were transferred to C57BL/6 mice before infection with X31-OVA. At 10dpi, OTI-TR2KO cells (magenta) were concentrated in the medulla region of the MLN, whereas GFP+ OTI-S4KO cells (green) were distributed in the cortex (right panel), including some OTI-S4KO cells that colocalized with HEVs (yellow) **(Fig 7A)**. Images of the lungs taken at 19dpi showed OTI-S4KO cells (green) in the airways (left panel) and OTI-TR2KO (magenta) near a large blood vessel (yellow) **(Fig 7B)**.

**Figure 7.**
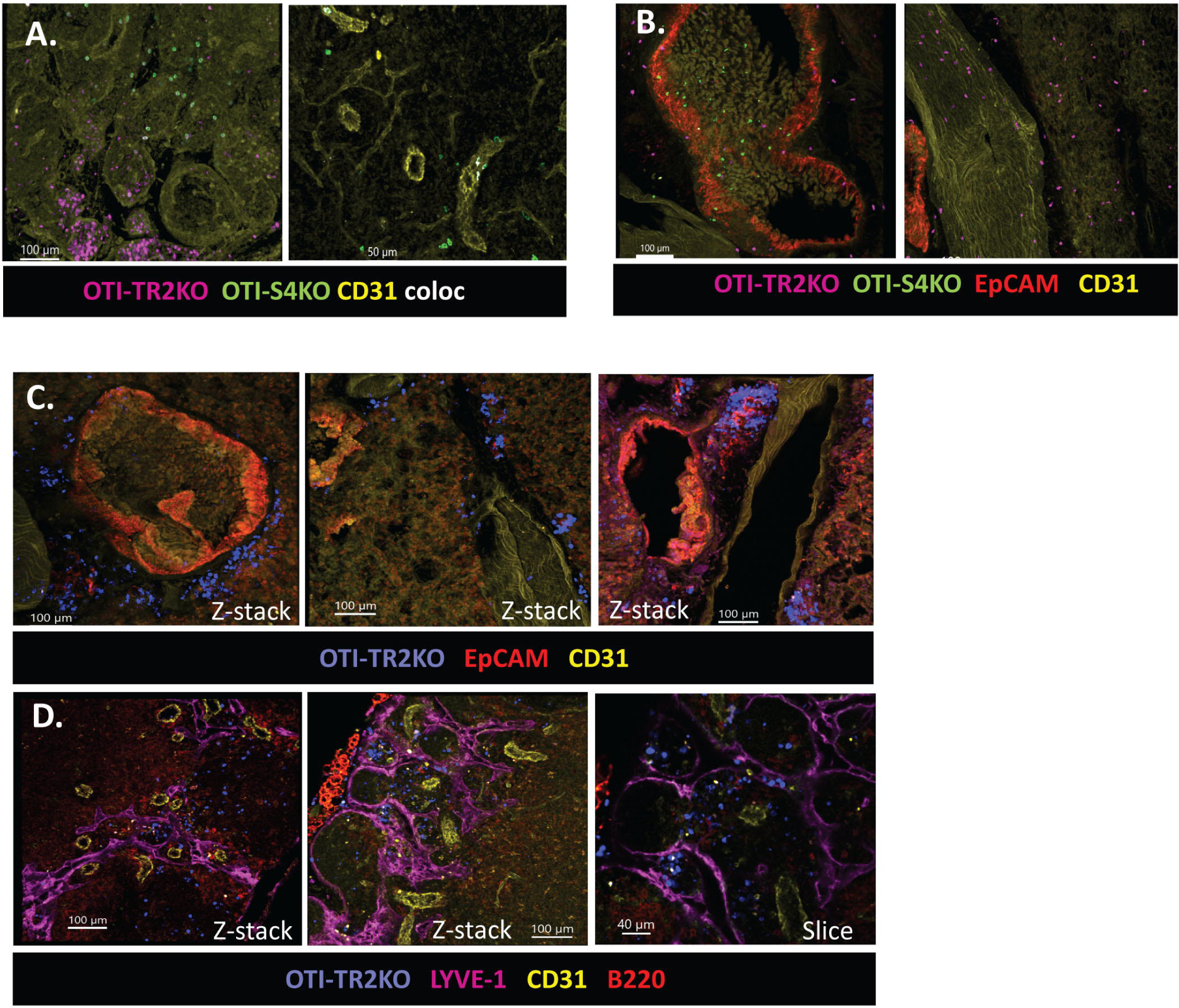
OTI-TR2KO and OTI-S4KO cells localize to different anatomical compartments. A&B) Mixed OTI-TR2KO and OTI-S4KO cells were transferred to C57BL/6 mice before infection with X31-OVA. A) OTI-TRKO cells (magenta) are clustered in the medulla region of the MLN at 10dpi. Interactions between OTI-S4KO cells (green) and HEVs (yellow) are show in white (right panel). B) OTI-S4KO cells (green) were located in the airways at 19dpi (left panel). OTI-TR2KO (magenta) were near a large blood vessel (yellow) - right panel. C&D) OTI-TR2KO cells were analyzed 21dpi with X31-OVA. C) OTI-TR2KO cells (blue) in the perivascular and peribronchial space of the lungs. D) OTI-TR2KO cells (blue) in the medulla region of the MLN.

To examine the MLN in more detail, we transferred OTI-TR2KO cells to C57BL/6 mice before infection with X31-OVA **(Fig 7C & 7D)**. At 21dpi, fixed sections of MLN were stained with antibodies specific for LYVE-1 (magenta) to visualize the medulla region of the lymph node. Some OTI-TR2KO cells (blue) were located in the perivascular and peribronchial space of the lungs **(Fig 7C)**, as well as the medulla region of the MLN **(Fig 7D)**. The images of the lungs are not entirely consistent with our prior study, where i.v. staining suggested that most KLRG1^+^ CTLs were inside the blood vessels (22). This discrepancy could reflect leakage of injected antibody from defuse lung tissue during IV staining.

## Discussion

In this study, confocal imaging has been used to explore how terminal differentiated CTLs distribute in the lungs during respiratory infection. We found KLRG1^+^ CTLs in the microvasculature and large blood vessels of the lungs, but not the lumen of the airways. In mixed transfer experiments the airways were preferentially populated with OTI-S4KO cells that expressed KLRG1 and CX_3_CR1 at reduced levels. Chemokine receptor CXCR6 has previously been implicated for a role in localization of T_RM_ cells into the airways(39). We transcriptome analysis to analyze gene expression during IAV infection and found that similar levels of CXCR6 expression in OTI-S4KO and OTI-Ctrl cells(17).

Our images showed N-cadherin expression on smooth muscle cells in the lungs and around HEVs in the MLN. We also found show that OTI-S4KO and OTI-TR2KO cells localize in different regions of the MLN. OTI-TR2KO cells expressed KLRG1 at high levels and were primarily located in the medulla region, whereas OTI-S4KO were found in cortex and interacted with HEV. Further work is required to further define the function of cadherin-binding proteins during T cell migration, determine the functional importance of interactions between KLRG1 and N-cadherin. These molecules may enhance retention of CTLs inside the blood vessels or adversely affect T cell survival in the periphery. These studies emphasize the importance of the inflammatory milieu during CD8 T cell differentiation and point to specific roles for KLRG1 and CD103 during tissue localization.

## Acknowledgments

We thank the histology core and the center for cell analysis and modeling (CCAM) at UCONN Health for assistance with this study.

## Material and methods

### Mice and reagents

Mice were bred and housed at the University of Connecticut Health Center in accordance with institutional guidelines. Experiments were performed in accordance with protocols approved by the UCONN Health Institutional Animal Care and Use Committee (IACUC). Mice that express Cre-recombinase under the control of the distal-Lck promoter (dLck) were used to generate mice with CD8 T cells that lack SMAD4 or TGFβ receptor II (TR2KO)(17). These mice were further crossed with OTI mice that express a transgenic antigen receptor specific for the SIINFEKL peptide presented on H-2Kb(40). In addition, OTI-Ctrl and OTI-S4KO mice were crossed with reporter mice that express GFP under the control of the fractalkine receptor (CX_3_CR1)(29). Virus stocks were grown in fertilized chicken eggs (Charles River) and stored as described previously. Between 8 to 20 weeks (wks) after birth, anesthetized mice were infected intranasally (i.n.) with either 2×10^3^ plaque forming units (PFU) X31-OVA, or 5×10^3^ colony-forming units (CFU) of recombinant Listeria monocytogenes expressing chicken ovalbumin (LM-OVA)(27).

### Adoptive cell transfer and sample preparation for flow cytometry

Naive CD8 T cells were isolated from SLO using Mojosort isolation kits (Biolegend, Dedham MA). Mice received 5×10^3^ congenically-marked donor cells by intravenous (i.v.) injection given 48 hours (hrs) before infection. To isolate CTLs for flow cytometry, chopped lung tissue was incubated at 37°C for 90 minutes (mins) in RPMI with 5% fetal bovine serum (FBS) and 150 U/ml collagenase (Life Technologies, Rockville, MD, USA). Nonadherent cells were enriched on Percoll density gradients (44/67%) spun at 1200g for 20 min. Lymphocytes were incubated with antibodies that block Fc-receptors (15 mins at RT). Antigen-experienced CD8 T cells were identified using high CD11a/CD44 expression and divided into subsets with antibodies specific for CD103, KLRG1, CD127 and CD62L. For intracellular staining, lymphocytes were analyzed using True Nuclear transcription factor buffer (Biolegend, Dedham-MA). Permeabilized cells were stained with antibodies specific for T-bet and EOMES. Data were analyzed using Flowjo. and Graphpad Prism software.

### Confocal microscopy

Infected tissues were fixed in 4% paraformaldehyde (PFA)/PBS (1hr at 4°C) and incubated with antibodies to block Fc-receptors (15 mins at 4°C). Tissues were stained with antibodies to CD45.1, EpCAM, E-cadherin, N-cadherin and CD31. Donor cells were detected with antibodies specific for CD45.1. Washed tissues were mounted on slides using Shandon Immu-Mount (Thermo Electron, Pittsburgh, PA, USA). Images were recorded using a Zeiss LSM880 confocal microscope with an inverted Axio Observer. Fluorescence was detected using: an argon laser for emissions at 458, 488, and 514 nm; a diode laser for emissions at 405 and 440 nm; a diode-pumped solid-state laser for emissions at 561 nm; and a HeNe laser for emissions at 633 nm. Images were analyzed using the colocalization function in Imaris suite software (Bitplane, South Windsor, CT, USA).

### Histology

Tissues were fixed in 4% PFA/PBS at 4°C for 24-48 hrs. Washed tissues were stored in 70% ethanol until processing. Tissues were embedded in paraffin blocks, sectioned, and stained with hematoxylin and eosin (H&E) by the Histology Core at the UCONN Health. Images were takes at 5X and 20X normal magnification.

